# Crystal structure of plant γ-glutamyl peptidase 1 with dual roles in sulfur metabolism and implications for oxidative stress regulation

**DOI:** 10.1101/2024.11.09.622773

**Authors:** Kosei Sone, Takehiro Ito, Chihaya Yamada, Toma Kashima, Akimasa Miyanaga, Naoko Ohkama-Ohtsu, Shinya Fushinobu

## Abstract

γ-Glutamyl peptidase 1 (GGP1) plays a dual role in primary and secondary sulfur metabolism in *Arabidopsis thaliana*. During glutathione (GSH) turnover, GGP1 hydrolyzes the isopeptide bond of GSH to degrade the tripeptide into Glu and Cys-Gly. During glucosinolate and camalexin biosynthesis, GGP1 processes GSH conjugates, which have a large substituent at the thiol side chain, by hydrolyzing the same isopeptide bond of γ-Glu. In the present study, we determined the crystal structures of the following GGP1 forms: ligand-free, Glu complex, covalent γ-Glu intermediate, and disulfide-linked S-S inactive forms. The intermediate structure, in which γ-Glu is covalently linked to the nucleophile C100, was trapped by mutating the catalytic His to Asn (H192N). In the Glu complex and γ-Glu intermediate structures, Glu bound to the S1 subsite is extensively recognized by several hydrogen bonds. The substrate recognition of the Cys-Gly moiety at the S1’ and S2’ subsites was revealed by modeling GSH in the active site. Mutational analysis indicated that R206 plays an important role in substrate binding by forming a salt bridge with Gly at the S2’ subsite. An open pocket is present beyond the thiol side chain of Cys in the S1’ subsite, which contributed to the dual activity of GGP1 toward GSH and GSH conjugates. The S-S inactive structure was obtained by soaking GGP1 crystals in Cys-Gly, and the catalytic cysteine (C100) partially formed a disulfide bond with a neighboring C154 residue. The partial inactivation of GGP1 in the presence of a pro-oxidant (Cys-Gly) has revealed its possible role in oxidative stress regulation in *Arabidopsis*.

## Introduction

Glutathione (GSH, γ-L-Glu-L-Cys-Gly) is a tripeptide containing an isopeptide bond between the γ-carboxy group of Glu and Cys (Supplementary Fig. S1). GSH plays various essential roles in plants, one of which is the maintenance of cellular redox homeostasis [1–3]. Two GSH molecules form a disulfide bond at their Cys residues to yield glutathione disulfide (GSSG). Moreover, mixed disulfides are formed with proteins or other chemical compounds in addition to GSH, and the thiol group of GSH is possibly oxidized to sulfenic, sulfinic, or sulfonic acid. These oxidation reactions consume excess reactive oxygen species (ROS) [4]. GSSG is reduced by glutathione reductase with NADPH to form two GSH molecules [5], and this reversible redox reaction alleviates oxidative stress [6]. Furthermore, GSH serves as a major sulfur repository. Because the thiol group of free Cys has higher reactivity than that of GSH, elevated intracellular Cys levels are highly toxic to organisms [7]. Thus, in plants, GSH serves as a storage and transport form of sulfur instead of Cys [8,9].

γ-Glutamyl peptidase 1 (GGP1, EC 3.4.19.16) is a cytosolic enzyme that hydrolyzes the isopeptide bond of GSH conjugates to Glu and Cys-Gly conjugates in the biosynthesis of glucosinolates and camalexin [10]. Among the five GGP genes in *Arabidopsis thaliana* (*GGP1*–*5*), *GGP1* and *GGP3* are highly expressed in all tissues and are involved in the biosynthesis of glucosinolates and camalexin through the processing of GSH conjugates and GSH-indole-3-acetonitrile (GS-IAN, Supplementary Fig. S1) [11]. We previously reported that GGP1, and possibly GGP3, are involved in GSH turnover [12]. *GGP1* is expressed under both sulfur-deficient and normal conditions and exhibits high GSH degradation activity to form Glu and Cys-Gly, the latter of which is then utilized for sulfur and protein metabolism [13]. Accordingly, GGP1 has dual roles in primary and secondary metabolisms through hydrolysis of the γ-Glu-Cys isopeptide bond in both GSH and GSH conjugates. Moreover, GGP1 is classified as part of the cysteine peptidase family C26 (γ-glutamyl hydrolase family) in the MEROPS database [14]. In this study, we report the crystal structures of four different forms of GGP1 in the active site, thereby providing structural insights into the dual activity of GGP1 and its possible regulation by redox reactions.

## Results and Discussion

### Overall structure

C-terminally His_6_-tagged recombinant protein of GGP1 was expressed in *Escherichia coli* and purified as a single band on SDS-PAGE. The crystal structure of ligand-free GGP1 was determined at 1.9 Å resolution (Table 1). The crystal belonged to space group *P*2_1_2_1_2_1_, and the asymmetric unit contained two GGP1 molecules. Molecular interface analysis using the PISA server [15] suggested that the protein is a monomer in solution. The final model contained the protein residues 4-246 in chain A and 4-250 in chain B. The overall structure of GGP1 comprises seven α-helices, two 3_10_ helices, and eleven β-strands, adopting a three-layer α/β/α sandwich fold (Fig. 1). The central β-sheet consists of six parallel strands (β2, β1, β3, β4, β11, and β9) and an antiparallel strand (β10). This β-sheet is sandwiched between three α-helices (α1, α2, and α7) and two β-strands (β6 and β7) on one side and two α-helices (α3 and α4), two 3_10_ helices (η1 and η2), and two β-strands (β5 and β8) on the other side. A two-helix bundle (α5 and α6) is extended from the core α/β/α fold. The catalytic triad of GGP1 comprises C100, H192, and E194. These residues are completely conserved in the C26 family and play essential roles in catalysis [16,17]. Therefore, the reaction mechanism of GGP1 was presumed to follow that of general cysteine peptidases (Supplementary Fig. S2). C100 is located in the loop between β4 and α4, while H192 and E194 are in the loop between β11 and α5 (Fig. 1). In the crystal structure of ligand-free GGP1, the distance between H192 and E194 is 2.7 Å, where a hydrogen bond is formed, and the distance between C100 and H192 is 3.6 Å (Fig. 2A). Although this distance is within the acceptable range for a hydrogen bond between the sulfur and nitrogen atoms, the relative orientation of the C100 side chain was not compatible with that of the hydrogen bond. This structural arrangement of the catalytic triad is common among C26 family peptidases [17].

**Fig. 1.**
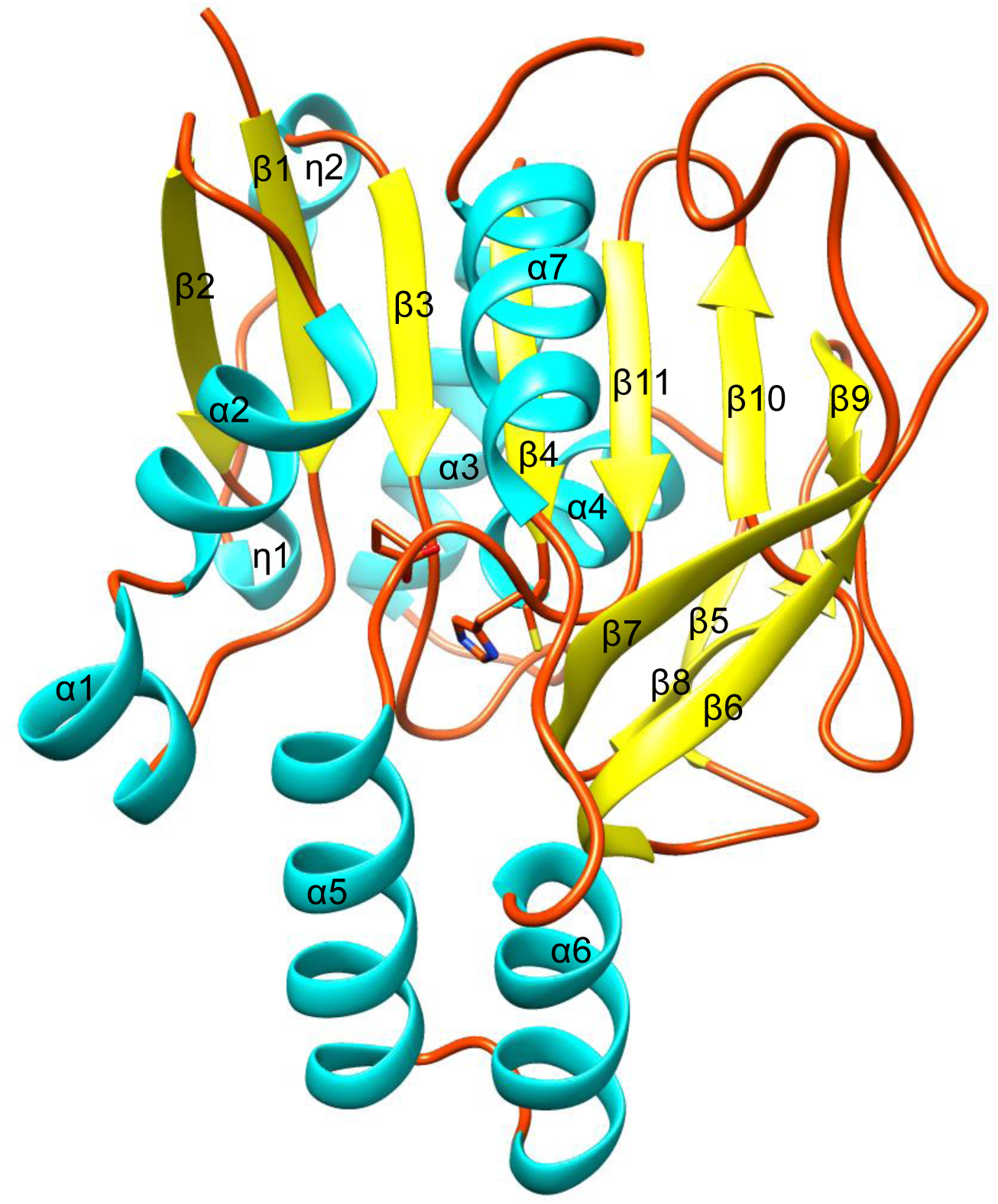
Overall structure of GGP1 in ligand-free form. The side chains of the catalytic triad (C100, H192, and E194) are shown as sticks.

**Fig. 2.**
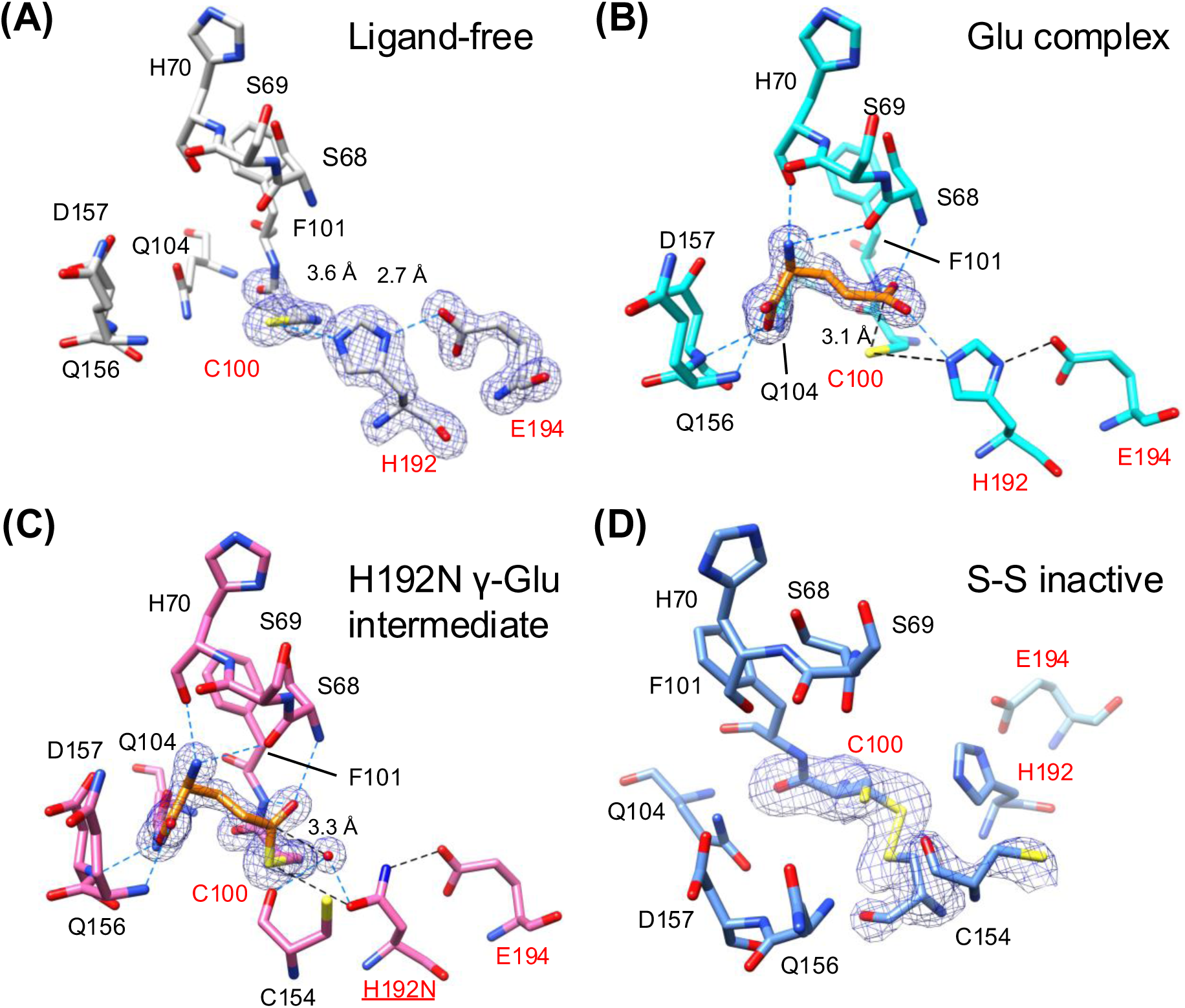
The active site of four GGP1 structures. (A) Ligand-free form, (B) Glu complex structure, (C) γ-Glu intermediate structure, and (D) S-S inactive structure. In (A), (B), and (C), polder maps at 6σ are shown. In (D), a polder map at 3.5σ is shown to indicate the two different conformations of C100 and C154. Hydrogen bonds with the ligand and within the catalytic triad are shown as blue and black dotted lines, respectively.

**Table 1.** Crystallographic data collection and refinement statistics.

| | Ligand-free | Glu complex | H192N γ-Glu intermediate | S-S inactive |
| --- | --- | --- | --- | --- |
| **Data collection** |  |  |  |  |
| Beamline | SPring-8 BL45XU | SPring-8 BL45XU | KEK PF-AR NE-3A | SLS PSI X06SA |
| Wavelength (Å) | 1.000000 | 1.000000 | 1.000000 | 1.000030 |
| Resolution range (Å) | 46.67–1.90 (1.94–1.90) | 46.63–1.74 (1.78–1.74) | 46.10–1.59 (1.62–1.59) | 46.51–2.04 (2.10–2.04) |
| Space group | $P2_12_12_1$ | $P2_12_12_1$ | $P2_12_12_1$ | $P2_12_12_1$ |
| Unit cell (Å) | $a = 72.542, b = 75.589, c = 103.137$ | $a = 72.289, b = 75.816, c = 102.772$ | $a = 71.508, b = 75.109, c = 101.206$ | $a = 72.354, b = 75.329, c = 102.583$ |
| Total reflections | 86197 (5576) | 111190 (6095) | 467799 (15838) | 241573 (19347) |
| Unique reflections | 45322 (2888) | 58231 (3148) | 71347 (2936) | 36242 (2787) |
| Multiplicity | 1.9 (1.9) | 1.6 (1.9) | 6.6 (5.4) | 6.7 (6.9) |
| Completeness (%) | 100.0 (100.0) | 100.0 (100.0) | 97.0 (82.7) | 100.0 (99.9) |
| Mean $I/\sigma(I)$ | 11.8 (1.6) | 15.3 (1.8) | 16.2 (1.8) | 13.5 (3.4) |
| $R_{\text{merge}}$ | 0.057 (0.467) | 0.038 (0.405) | 0.060 (0.603) | 0.096 (0.569) |
| $R_{\text{pim}}$ | 0.057 (0.467) | 0.038 (0.405) | 0.025 (0.279) | 0.040 (0.232) |
| $CC_{1/2}$ | 0.996 (0.663) | 0.999 (0.645) | 0.996 (0.791) | 0.998 (0.895) |
| Wilson B-factor (Å<sup>2</sup>) | 11.29 | 15.28 | 13.89 | 22.91 |
| **Refinement** |  |  |  |  |
| Resolution range (Å) | 46.72–1.90 | 46.67–1.74 | 46.15–1.59 | 46.55–2.04 |
| $R_{\text{work}}$ | 0.170 | 0.155 | 0.166 | 0.177 |
| $R_{\text{free}}$ | 0.219 | 0.195 | 0.200 | 0.233 |
| Number of atoms |  |  |  |  |
| Amino acids | 4018 | 3982 | 4047 | 4019 |
| Ligands | 14 | 18 | 32 | 18 |
| Waters | 347 | 466 | 448 | 126 |
| RMS deviations |  |  |  |  |
| Bonds lengths (Å) | 0.0149 | 0.0148 | 0.0102 | 0.0150 |
| Bond angles (°) | 2.441 | 2.340 | 1.843 | 2.554 |
| Ramachandran plot (%) |  |  |  |  |
| Favored | 96.93 | 97.14 | 97.58 | 96.92 |
| Allowed | 3.07 | 2.86 | 2.42 | 3.08 |
| Outlier | 0.00 | 0.00 | 0.00 | 0.00 |
| Average B-factor (Å<sup>2</sup>) |  |  |  |  |
| Amino acids | 23.91 | 22.29 | 21.40 | 30.59 |
| Ligands | 30.54 | 41.66 | 44.68 | 38.34 |
| Waters | 30.12 | 30.45 | 30.88 | 27.88 |
| PDB ID | 9K7J | 9K7I | 9K7L | 9K7K |

\*Values in parentheses represent the highest resolution shell.

### Ligand complex structures

The GGP1-Glu complex structure was determined at a resolution of 1.74 Å using a co-crystallization method (Table 1). The Glu complex crystal was isomorphous with the ligand-free crystal, and their main chain structures were similar. Their root mean square deviations (RMSD) for the Cα atoms was 0.24 Å. The electron density of Glu was clearly observed in the active site, and the bound Glu forms numerous hydrophilic interactions with the protein (Fig. 2B). The γ-carboxy group of Glu is located near the nucleophilic C100 residue, representing the product complex after hydrolysis of the thioester intermediate in the reaction cycle (Supplementary Fig. S2). According to the subsite nomenclature of peptidases [18], γ-Glu, Cys, and Gly of GSH correspond to P1, P1’, and P2’, and bind to the S1, S1’, and S2’ subsites, respectively. Therefore, this Glu molecule (P1) was bound to the S1 subsite of GGP1. The α-carboxy group of Glu forms hydrogen bonds with the main chain amides of D157 and Q156 and the side chain amide of Q104, whereas the amino group forms bifurcated hydrogen bonds with the main chain carbonyl groups of S68 and H70. The γ-carboxy group is hydrogen-bonded with the main chain amides of S68 and F101 and the side chain of H192. The Glu complex structure indicates that the main chain amides of S68 and F101 form the oxyanion hole, which is crucial for the catalytic function of serine and cysteine hydrolases and plays a pivotal role in stabilizing the tetrahedral intermediate during the reaction [19].

A previous study showed that a His-to-Asn mutant of the catalytic triad of the small (glutaminase) subunit of carbamoyl phosphate synthetase trapped a covalent intermediate structure with nucleophilic Cys [20]. Therefore, we co-crystallized the H192N mutant of GGP1 with GSH and successfully obtained a high-resolution (1.59 Å) intermediate structure in which C100 and γ-carboxy of Glu formed a covalent thioester bond (Table 1 and Fig. 2C). The electron density of the thioester bond was evident, and this structure mimicked the covalent intermediate state of cysteine peptidase (Supplementary Fig. S2, state 3). The Cα RMSD between the γ-Glu intermediate and ligand-free structures is 0.32 Å. H192 in the catalytic triad facilitates the deprotonation of nucleophilic water during the thioester hydrolysis step (Supplementary Fig. S2, state 5) as well as the nucleophilic attack of C100 (state 2). Therefore, the H192N mutation blocked thioester release and allowed the trapping of the covalent intermediate structure. Although the positions of the catalytic triad in the covalent complex structure are almost identical to those in the Glu complex structure, the thioester bond formation induced a slight movement of the γ-carboxyl group of Glu toward C100 (Supplementary Fig. S3). Interestingly, a prominent electron density peak for a water molecule was observed near the thioester carbon atom (Fig. 2C). This water is at a distance of 3.3 Å from the Cδ atom of Glu and forms hydrogen bonds with the N192 side chain and the main chain carbonyl of C154. We hypothesize that this water represents the position of nucleophilic water; however, the amide side chain of N192 could not facilitate hydrolysis of the thioester bond.

### Comparison with structural homologs

Structural homologs of GGP1 were searched using the Dali server [21]. Table 2 lists the top nine structures with the highest Z-scores. Three of these structures (PDB IDs: 3M3P, 3L7N, and 2H2W) remain unpublished, and one is a putative class I glutamine amidotransferase (GATase) from *Thermotoga maritima* (PDB ID: 1O1Y) [22]. The structural homologs of GGP1 with known enzyme activities include the glutaminase subunit of GATase from *Mycolicibacterium smegmatis* (MsGATase) [23], the TrpG (glutaminase) subunit of anthranilate synthase (EC 4.1.3.27) from *Saccharolobus solfataricus* (SsTrpG) and *Serratia marcescens* (SmTrpG) [24,25], and homoserine *O*-succinyltransferase (E.C 2.3.1.46) from *E. coli* and *T. maritima* [26,27]. Putative *T. maritima* GATase, SsTrpG, and SmTrpG form part of the Class I glutamine amidotransferases family, according to the Structural Classification Of Proteins-extended (SCOPe) classification[28]. MsGATase and the two TrpG proteins exhibit glutaminase activity that hydrolyzes Gln to Glu and NH_3_ (EC 3.5.1.35), and their structures have folds similar to that of GGP1 (Supplementary Fig. S4).

**Table 2.**
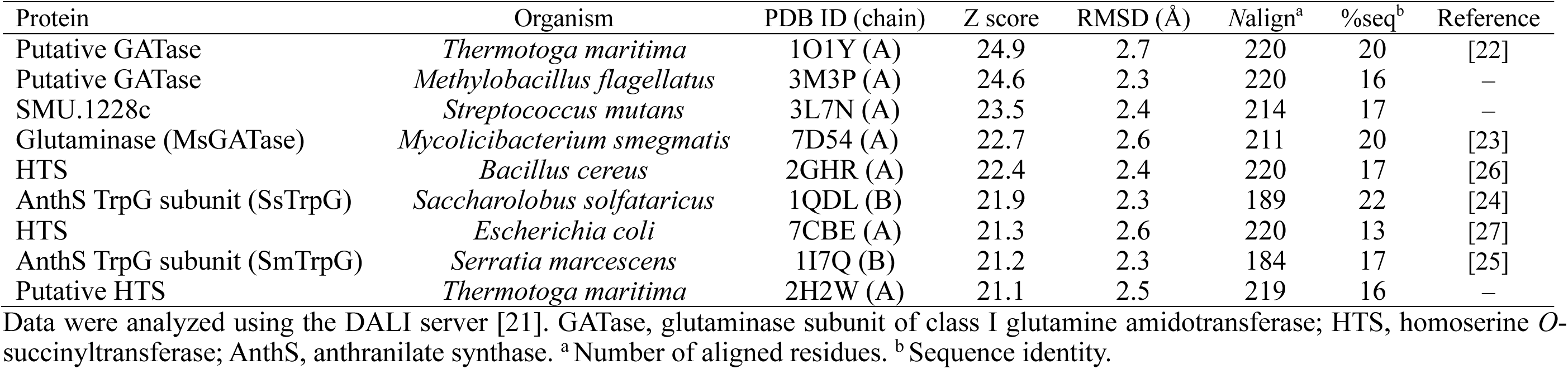
Structural similarity search results.

| Protein | Organism | PDB ID (chain) | Z score | RMSD (Å) | Nalign<sup>a</sup> | %seq<sup>b</sup> | Reference |
| --- | --- | --- | --- | --- | --- | --- | --- |
| Putative GATase | *Thermotoga maritima* | 1O1Y (A) | 24.9 | 2.7 | 220 | 20 | [22] |
| Putative GATase | *Methylobacillus flagellatus* | 3M3P (A) | 24.6 | 2.3 | 220 | 16 | – |
| SMU.1228c | *Streptococcus mutans* | 3L7N (A) | 23.5 | 2.4 | 214 | 17 | – |
| Glutaminase (MsGATase) | *Mycolicibacterium smegmatis* | 7D54 (A) | 22.7 | 2.6 | 211 | 20 | [23] |
| HTS | *Bacillus cereus* | 2GHR (A) | 22.4 | 2.4 | 220 | 17 | [26] |
| AnthS TrpG subunit (SsTrpG) | *Saccharolobus solfataricus* | 1QDL (B) | 21.9 | 2.3 | 189 | 22 | [24] |
| HTS | *Escherichia coli* | 7CBE (A) | 21.3 | 2.6 | 220 | 13 | [27] |
| AnthS TrpG subunit (SmTrpG) | *Serratia marcescens* | 1I7Q (B) | 21.2 | 2.3 | 184 | 17 | [25] |
| Putative HTS | *Thermotoga maritima* | 2H2W (A) | 21.1 | 2.5 | 219 | 16 | – |

Data were analyzed using the DALI server [21]. GATase, glutaminase subunit of class I glutamine amidotransferase; HTS, homoserine *O*-succinyltransferase; AnthS, anthranilate synthase. <sup>a</sup> Number of aligned residues. <sup>b</sup> Sequence identity.

The crystal structure of MsGATase complexed with Gln was obtained using a co-crystallization method [23]. Fig. 3A shows a superimposition of the Glu complex structure of GGP1 and the Gln complex structure of MsGATase (chain B). Although chain A of MsGATase bound to Gln in a catalytically irrelevant reverse orientation (not shown), Gln (magenta in Fig. 3A) in chain B (green in Fig. 3A) was bound in an apparently correct manner, placing the side chain amide nitrogen near the catalytic Cys. Although it is unclear why the Gln substrate was not hydrolyzed during the crystal growth of MsGATase, the catalytic residue (Cys99) was oxidized to cysteine sulfonate, and the bound ligand was assigned as a Gln molecule (not Glu). The superimposition of GGP1 and MsGATase indicated that their ligand-binding modes were notably different (Fig. 3A). The α-carboxy and amino groups of the Glu molecule in the S1 subsite of GGP1 are recognized by multiple residues, whereas those groups of Gln in MsGATase have no evident interactions with the protein. Considering that the residues recognizing the main chain groups of Glu in GGP1 (S68, H70, Q104, Q156, and D157) are highly conserved in MsGATase, they may share the same subsite S1 architecture, and the binding mode of Gln in chain B of MsGATase is probably incorrect.

**Fig. 3.**
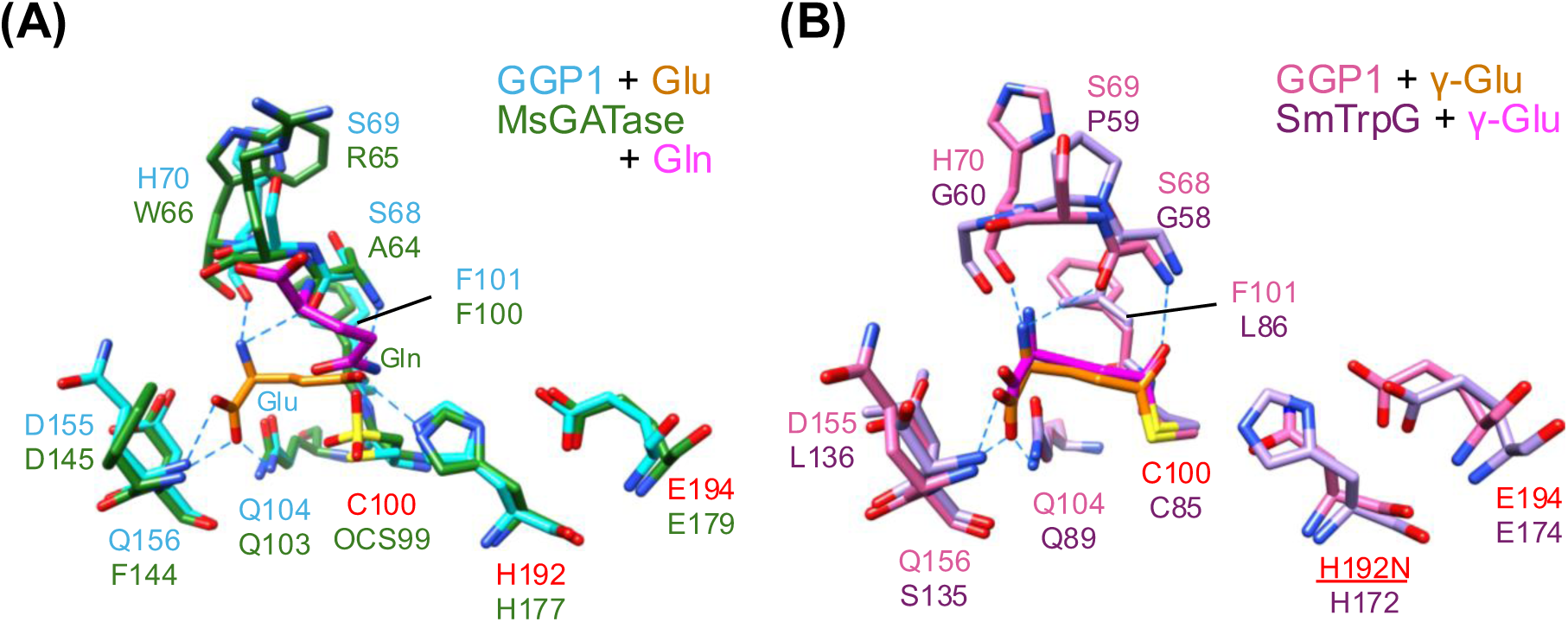
Comparison with glutaminases. (A) Superimposition of the Glu complex structure of GGP1 (protein in cyan and Glu in orange) and glutaminase subunit of glutamine amidotransferase from *M. smegmatis* (MsGATase, PDB ID: 7D54, chain B in green) complexed with Gln (magenta). (B) Superimposition of the γ-Glu intermediate structure of GGP1 (protein in pink and Glu moiety in orange) and TrpG (glutaminase) subunit of anthranilate synthase from *S. marcescens* (SmTrpG, protein in purple and Glu moiety in magenta).

Spraggon et al. successfully obtained a glutamyl thioester intermediate of SmTrpG with no mutations in the catalytic triad [25]. When the γ-Glu intermediate structure of GGP1 was overlayed on the γ-Glu intermediate structure of SmTrpG (Fig. 3B), the positions of the catalytic triad and Glu were nearly identical, indicating that the substrate recognition at the S1 subsite is conserved in these enzymes. The Class I glutamine amidotransferases family in SCOPe and the C26 γ-glutamyl hydrolase family in MEROPS generally have specificity for the isopeptide bond cleavage of P1 Glu, and this specificity is common to glutaminase that hydrolyzes the side chain amide nitrogen of Gln. Therefore, we concluded that the S1 subsite recognition architecture is basically conserved in these enzyme families.

### S-S inactive structure

Although analysis of the Glu complex and γ-Glu intermediate structures helped elucidate the S1 subsite (γ-Glu) recognition architecture of GGP1, the substrate recognition at subsites S1’ and S2’ for the Cys-Gly dipeptide remained unknown. We soaked wild-type GGP1 crystals in a solution containing Cys-Gly dipeptide and collected a diffraction dataset up to 2.04 Å resolution (Table 1). Unexpectedly, the electron density map showed that the S1’ and S2’ subsites were not occupied, and C100 partially formed a disulfide bond with C154 in the next β-strand (β7) (Fig. 2D). This is an inactive state because C100 is the catalytic nucleophile residue. We also observed a disulfide-free state in the crystal, and the occupancies of the S-S inactive and disulfide-free states were refined to 0.7 and 0.3, respectively, based on the electron density map (Supplementary Fig. S5). The ligand-free and S-S inactive structures are nearly the same except for the disulfide bond in the active site; the Cα RMSD between them is 0.32 Å. Considering that the Cys-Gly dipeptide is a pro-oxidant that acts as an iron reductant as well as a lipid peroxidation inducer [29], soaking crystals in the dipeptide-containing solution may have prompted the oxidization of two neighboring cysteine residues. In plant cells, GSH functions as an antioxidant by consuming excess ROS in the cytosol through reversible redox reactions between GSH and GSSG [1]. We hypothesized that the disulfide bond involving the catalytic C100 residue is formed under oxidative stress and that the loss of GGP1 function prevents the degradation of the antioxidant GSH. Fig. 4 shows the partial amino acid sequence alignment of 30 plant GGP1 homologs with the lowest protein BLAST e-values (<1×10^™162^), indicating that C154 is not conserved among plants. Several response pathways for oxidative stress in *Arabidopsis* have been reported [11,30–32], and numerous factors are involved in the regulation of GSH supply under oxidative stress. The GGP1 inactivation by disulfide bond formation may be one such factor in the regulation of oxidative stress in *Arabidopsis*.

**Fig. 4.**
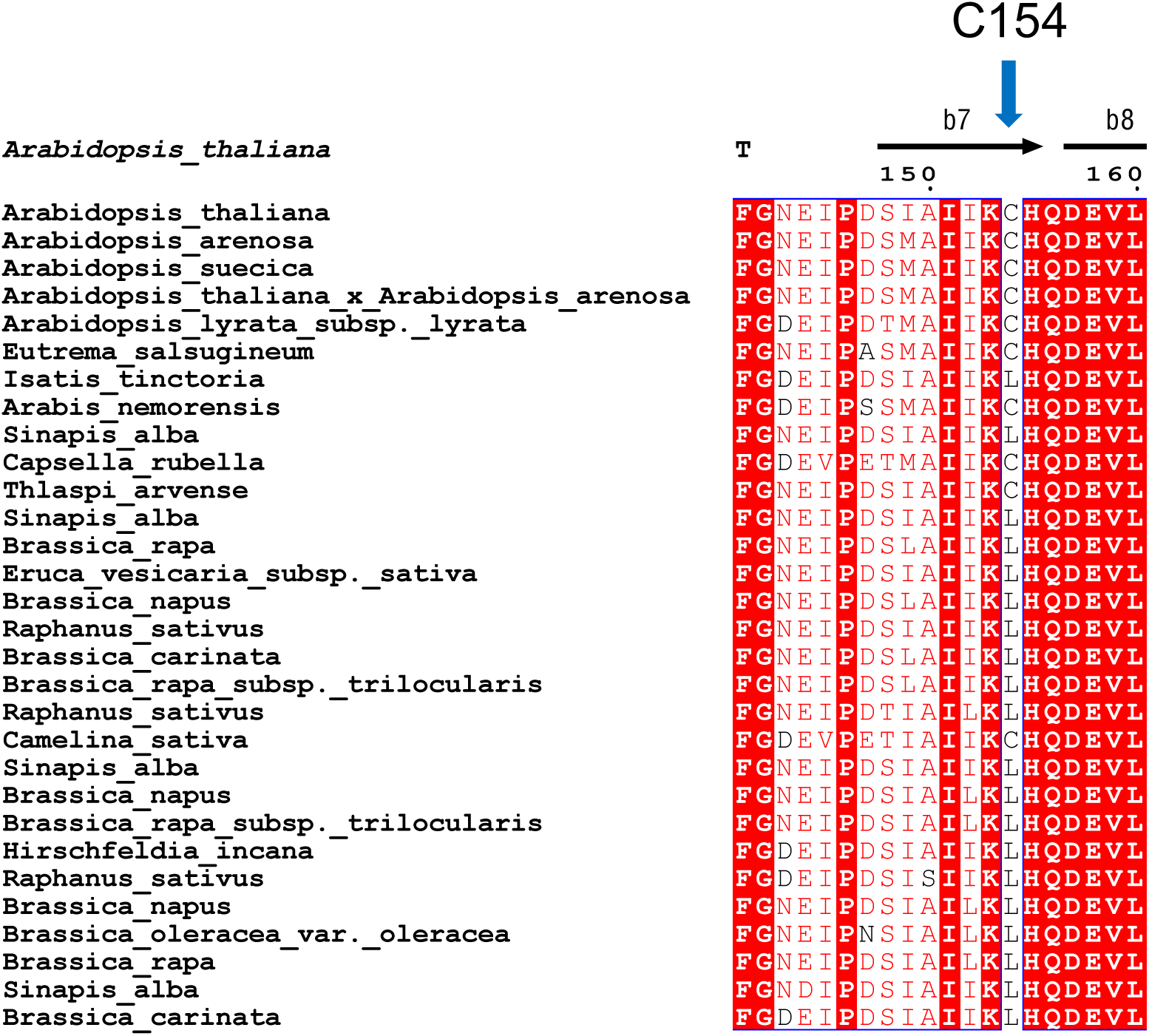
Partial amino acid sequence alignment of plant GGP1 homologs around C154.

### Substrate modeling and implications for dual physiological roles

A notable physiological feature of GGP1 is its dual role in primary (GSH degradation) and secondary (processing GSH conjugates) sulfur metabolism [12]. In the biosynthetic pathways of glucosinolates and camalexin, GGP1 cleaves the γ-Glu isopeptide bond of GSH conjugates or GS-IAN by accommodating a large thiol-linked substituent at P1’ Cys [11]. In a previous study, to examine the ability of GGP1 to process GSH and its conjugates, we modeled GSH in a predicted structure of GGP1 using AlphaFold [12]. In the present study, we constructed a more reliable GSH model in the active site of GGP1 based on the Glu complex and γ-Glu intermediate structures. The modeled GSH molecule forms extensive hydrogen bond interactions with GGP1, primarily at P1 γ-Glu (Fig. 5A). For the main chain of P1’ Cys, the amide nitrogen forms a hydrogen bond with the side chain of H192, and the carbonyl oxygen forms hydrogen bonds with the side chains of C154 and Y195. We previously suggested that R206 may form a salt bridge with the carboxylate group of P2’ Gly [12]. However, the R206 side chain was flipped away from the active site in the crystal structure. A surface representation of GGP1 with the modeled GSH (Fig. 5B) shows that the thiol side chain of P1’ Cys is pointed to an open pocket that can accommodate a large group of GSH conjugates or GS-IAN. The pocket comprises C154, Q156, and Y212, and these residues may affect the substrate specificity for GSH conjugates. GGP1 has a higher affinity for *S*-[(*Z*)-phenylacetohydroximoyl]-_L_-glutathione (GS-B, Supplementary Fig. S1) than for GSH [10,12]. This can be explained by a possible π-π stacking between Y212 and the phenyl group of GS-B.

**Fig. 5.**
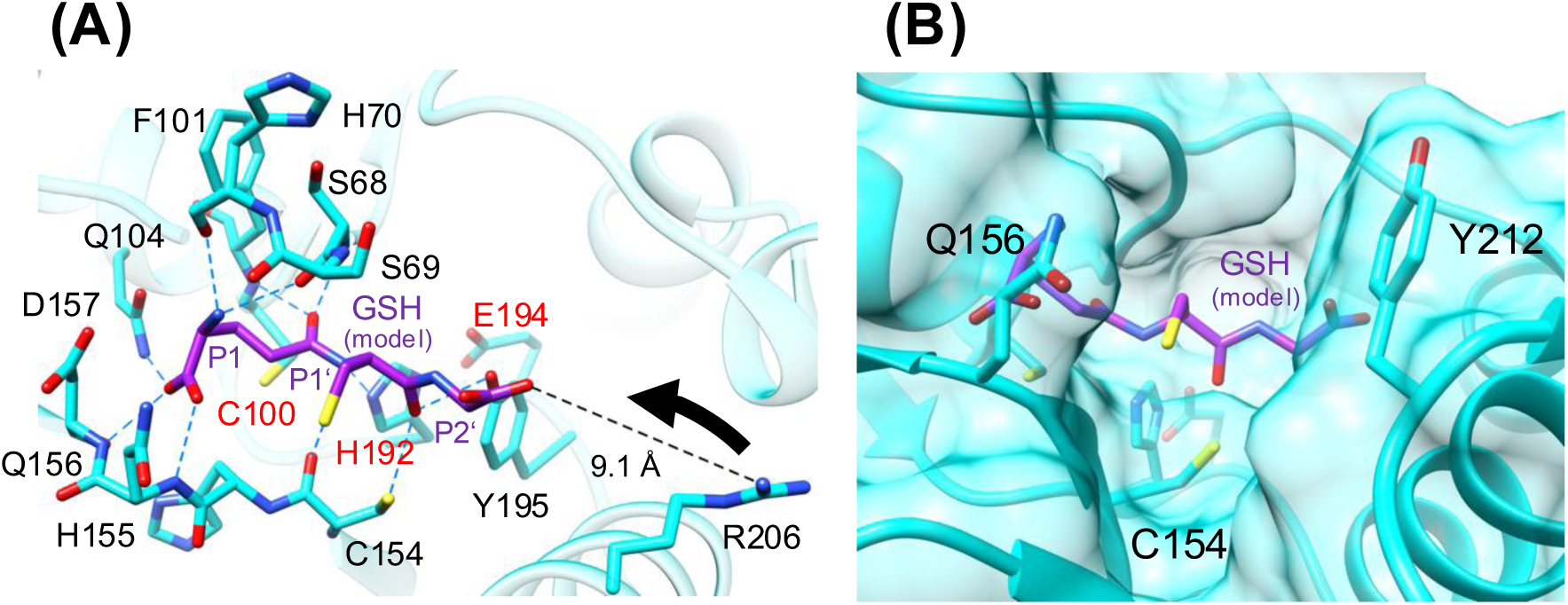
The active site of GGP1 with modeled GSH. (A) The active site interactions of GGP1 (cyan) with modeled GSH (purple). Hydrogen bonds are shown as blue dotted lines. The distance between the R206 side chain and the carboxy group of P2’ Gly is shown as a black dotted line. (B) The molecular surface of the active site of GGP (cyan) with modeled GSH (purple). The side chains of the catalytic triad residues and residues forming a pocket that accommodates the substituents of GSH conjugates (C154, Q156, and Y202) are shown.

### Mutational analysis

To verify the substrate interactions predicted by the substrate modeling, a site-directed mutational analysis of GGP1 was conducted. First, we optimized the activity assay condition. The *K*_m_ values of recombinant GGP1 toward GSH and a GSH conjugate (GS-B) were 5.0 mM and 37 μM, respectively [10,12]. In this study, we reexamined the buffer system to ensure that the pH did not change at high substrate concentrations. As shown in Fig. 6, the initial velocity did not saturate at 12.5 mM GSH in 200 mM HEPES-NaOH buffer (pH 7.5) and 200 mM NaCl. We determined that the pH of the previous assay condition (50 mM Tris-HCl buffer adjusted to pH 8.0) decreased at high GSH concentrations (>7.5 mM), and the activity also decreased at high substrate concentrations. Although we could not determine the *K*_m_ value of GGP1 toward GSH, the *k*_cat_/*K*_m_ value calculated from the slope of the plot was 36.1 s^-1^M^-1^ (Fig. 6). Considering that the cytosolic GSH concentration in *Arabidopsis* is 1–5 mM [2], the *K*_m_ value of GGP1 for GSH is far higher than its concentration in the cytosol.

**Fig. 6.**
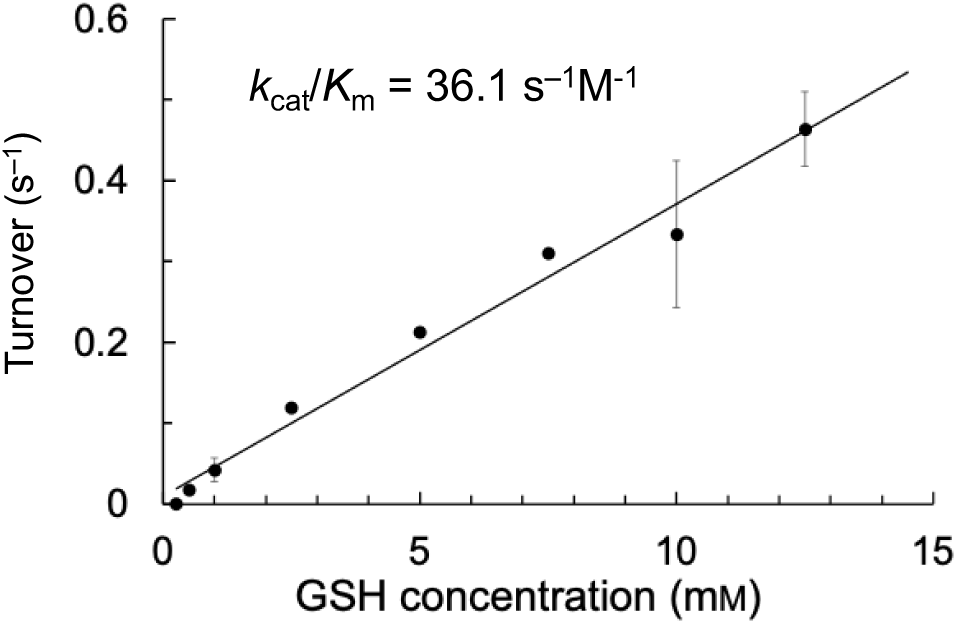
*S*-*v* plot of GGP1 activity toward GSH.

Single-site substitution mutants (C100S, H192N, C154S, C154A, and R206A) were constructed. All the mutants were expressed in *E. coli* and the activity of the purified enzyme was measured (Table 3). The activities of the two catalytic triad mutants, C100S and H192N, were not detected. The significant activity decrease of R206A indicates that R206 recognizes P2’ Gly and is essential for GSH degradation, whereas its side chain was flipped away from the active site in the crystal structure without Cys-Gly (Fig. 5A). We hypothesize an induced-fit motion of R206, in which the side chain moves toward P2’ Gly on the binding of the substrate. The importance of R206 is supported by the high degree of conservation among plant GGP1 homologs (Supplementary Fig. S6). C154 formed a disulfide bond with C100 in the S-S inactive structure (Fig. 2D) and was predicted to form a hydrogen bond with the main chain carboxy group of P1’ Cys (Fig. 5A). C154A retained 45% activity, whereas the activity of C154S was reduced to <5% (Table 3). In the circular dichroism (CD) spectra of the mutants, C100S retained a positive peak at <200 nm, whereas the peaks of the other mutants decreased (Supplementary Fig. S7). For the negative signal around 220 nm, which indicated helix content, the mutants showed a decrease in C100S, H192N, C154A/S, and R206A. Secondary structure content analysis showed that the mutants had decreased helical content and increased turn content (Supplementary Table S1). The decrease in helix content was particularly noticeable in C154S and R206A, suggesting that C154 and R206 also contributed to protein structure stabilization. The decrease in the activities of these mutants may be a result of the loss of substrate interactions as well as protein instability.

**Table 3.** Activity of site-directed mutants.

| Mutant | Activity (%) |
| --- | --- |
| C109S | ND |
| H192N | ND |
| C154S | 4.9 ± 1.7 |
| C154A | 45 ± 3 |
| R206A | 4.4 ± 1.0 |

Relative activity compared with that of the wild-type GGP1 ( $100 \pm 2.7\%$ ). ND, not detected (<1%). Activity toward 5 mM GSH at pH 7.5 and 37 °C was measured. All experiments were performed in triplicate ( $n = 3$ ) and expressed as mean ± standard error.

## Conclusions

In this study, we determined the high-resolution crystal structures of GGP1 from *A. thaliana*, which is categorized as part of the C26 γ-glutamyl hydrolase family of the MEROPS database. To the best of our knowledge, this is the first three-dimensional structure of a plant enzyme belonging to the C26 family to be determined. The Glu complex and γ-Glu intermediate structures indicate that GGP1 shares a common catalytic mechanism for cysteine hydrolases that involves nucleophilic attack by the thiol side chain of cysteine, formation of an oxyanion hole by the main chain amides, and the presence of nucleophilic water near the catalytic histidine. Moreover, the elaborate P1 γ-Glu recognition at the S1 subsite, which may be common in γ-glutamyl hydrolase family enzymes and glutaminases, was elucidated. These complex structures allowed us to model the GSH molecule, including the P1’-P2’ Cys-Gly moiety. The S-S inactive form indicates a possible physiological role of GGP1 in oxidative stress of *Arabidopsis* species. The structural basis for the dual physiological roles of GGP1 in primary and secondary metabolism was elucidated from the modeled structure with GSH in the active site. The thiol side chain of P1’ Cys was shown to be in an open pocket that accommodates a large substituent group, thereby explaining its activity toward GSH and its conjugates. The substrate specificity and catalytic efficiency of GGP1 can be modified for a particular GSH conjugate by mutating the pocket-forming residues. Because specific glucosinolates have been reported to exhibit chemopreventive actions that decrease cancer risk [33,34], engineering GGP1 may lead to the establishment of a production platform for glucosinolates in a controlled manner.

## Methods

### Recombinant protein production and purification

From the previously constructed expression plasmid[12], an unnecessary 18-amino-acid sequence (LESTSLYKKAGSAAAPFT) derived from pDEST17 and the Gateway entry vector pENTR/D-TOPO (Invitrogen, Carlsbad, CA, USA) was removed using PCR with the KOD One PCR Master Mix (TOYOBO Co., Ltd., Osaka, Japan) and the primers listed in Supplementary Table S2. The resulting plasmid contained a short His_6_-tag (MSYYHHHHHH) at the N-terminus of GGP1. *E. coli* BL21 (DE3) cells harboring the expression plasmid were cultured at 37 °C in lysogeny broth medium (1% tryptone, 0.5% yeast extract, and 1% NaCl) with 34 µg/mL ampicillin until the OD_600_ reached 0.5. Subsequently, overexpression was induced by adding 0.1 mM isopropyl β-D-thiogalactopyranoside and continued for 20 h at 16 °C. The cells were harvested through centrifugation at 8000 × *g* and resuspended in 50 mM HEPES-NaOH (pH 7.5) and 200 mM NaCl. The cells were disrupted using an ultrasonic homogenizer (Branson Sonifier250D; Branson Ultrasonics Division of Emerson Japan, Kanagawa, Japan), and the supernatant was collected through centrifugation at 15000 × *g* and filtered using Minisart Hydrophilic 0.45-µm filters (Sartorius Stedim Biotech, Göttingen, Germany). The crude enzyme solution was purified using Ni-affinity chromatography (cOmplete His-Tag; Roche Diagnostic GmbH, Mannheim, Germany). Solutions of 5 and 250 mM imidazole in 50 mM HEPES-NaOH (pH 7.5) and 200 mM NaCl were used as wash and elution buffers, respectively. The eluted fraction was concentrated using an ultrafiltration centrifugal membrane unit (Amicon Ultra 15 MWCO 10 kDa; Millipore, Billerica, MA, USA) and loaded onto a gel filtration chromatography column (HiLoad 16/60 Superdex 200 pg; Cytiva, Marlborough, MA, USA) equilibrated with 20 mM HEPES-NaOH (pH 7.5) and 200 mM NaCl. Protein concentrations were determined using the bicinchoninic acid (BCA) protein assay kit (Thermo Fisher Scientific, Waltham, MA, USA) or absorbance at 280 nm. The theoretical extinction coefficient was calculated from the amino acid sequence using the ProtParam tool in Expasy (https://web.expasy.org/protparam/). The purity of the enzymes was confirmed at each purification step using SDS-PAGE.

### Crystallography

Solutions of GGP1 were desalted before crystallization. Crystals were grown at 4 °C using the sitting drop vapor diffusion method. A protein solution (0.5–0.6 µL) containing 4–7 mg/mL protein was mixed with a reservoir solution (0.4–0.5 µL) containing 0.1 M Na-acetate (pH 4.0–4.75), 1.20–1.32 M K_2_HPO_4_, and 0.80–0.88 M NaH_2_PO_4_. Crystals of the Glu complex were prepared through co-crystallization with 7.5 mM Glu. For the γ-Glu intermediate, 5.0 mM GSH was co-crystallized. To prepare S-S inactive crystals, GGP1 crystals were soaked in 100 mM Cys-Gly dipeptide for 5 s. The crystals were cryoprotected in a reservoir solution supplemented with 20% glycerol or 20% PEG200 and flash-frozen in liquid nitrogen. X-ray diffraction data were collected at the beamlines of the Photon Factory of the High Energy Accelerator Research Organization (KEK, Tsukuba, Japan), the Swiss Light Source (SLS) of the Paul Scherrer Institut (PSI, Villigen, Switzerland), and SPring-8 (Hyogo, Japan). For the diffraction datasets collected at SPring-8, initial data processing was performed using KAMO [35] based on XDS [36]and DIALS [37]. Only XDS [36] was used for the other datasets. Data reduction to proper resolution was performed using Aimless software[38]. The initial structures were solved using the molecular replacement method implemented in PHASER [39] using a predicted structure provided by ColabFold[40]. Manual model building and refinement were performed using Coot [41] and Refmac5 [42]. Polder maps were prepared using PHENIX software[43]. Molecular graphic images were prepared using Chimera [44] and Chimera X software[45].

### Construction of mutants and enzyme assay

The KOD One PCR Master Mix (TOYOBO Co., Ltd.) was used for site-directed mutagenesis. The primers used for each mutant are listed in Supplementary Table S2. The reaction mixture for activity measurements (100 µL) contained 2.5 µg of the wild-type or mutant GGP1, 0.25–12.5 mM GSH, 200 mM HEPES-NaOH (pH 7.5), and 200 mM NaCl. To measure the activity of the mutants, 5 mM of GSH was used. The reaction mixtures were incubated at 37 °C and the reaction was terminated by heating at 95 °C. The solutions were centrifuged at 15000 × *g* for 15 min at 4 °C. The released Cys-Gly in the supernatants was quantified using HPLC after derivatization with monobromobimane, as previously described [12].

### Measurement of CD spectra

CD spectra were measured using a J-820 spectropolarimeter (JASCO Corp., Hachioji, Tokyo, Japan). Solutions of wild-type GGP1, and C154S and C154A mutants were replaced with 10 mM NaHPO_4_ (pH 7.5) using a PD-10 column (Sigma-Aldrich, St. Louis, MI, USA). The averages of the four scans are presented as the molar ellipticity per residue. The secondary structure was estimated using the BeStSel server[46].

## Acknowledgments

The authors thank Dr. Takatoshi Arakawa for the valuable discussions. We also thank the staff of KEK-PF, SLS at PSI, and SPring-8 for the X-ray data collection. This research was supported by JSPS-KAKENHI (19H02859 to NOO, and 19H00929 and 23H00322 to SF) and partly by the Research Support Project for Life Science and Drug Discovery (Basis for Supporting Innovative Drug Discovery and Life Science Research (BINDS)) from AMED under Grant Number JP22ama121001.

## Author contributions

NOO and SF conceived and supervised the study; CY, TA, TK, AM, and SF planned the experiments; KS and TI performed the experiments; KS performed the protein crystallography; TI performed the biochemical experiments and; KS and SF wrote the manuscript. All authors reviewed the final version of the manuscript.

## Data availability statement

Atomic coordinates and structure factors of the crystal structures have been deposited in the Protein Data Bank under accession IDs 9K7I, 9K7J, 9K7K, and 9K7L. The source data are provided with this paper.

## Abbreviations

GSH: glutathione
GSSG: glutathione disulfide
GGP: γ-glutamyl peptidase
GS-IAN: GSH-indole-3-acetonitrile
RMSD: root mean square deviation
GATase: glutamine amidotransferase
MsGATase: *Mycolicibacterium smegmatis* GATase
SsTrpG: TrpG subunit of *Saccharolobus solfataricus* anthranilate synthase
SmTrpG: TrpG subunit of *Serratia marcescens* anthranilate synthase
GS-B: *S*-[(*Z*)-phenylacetohydroximoyl]-L-glutathione
CD: circular dichroism
SCOPe: Structural Classification of Proteins-extended
ROS: reactive oxygen species;
PDB: Protein Data Bank

## Notes

**Conflict of interest:** The authors declare no conflict of interest.

### Competing Interest Statement

The authors have declared no competing interest.

## References

1 Foyer CH & Noctor G (2011) Ascorbate and Glutathione: The Heart of the Redox Hub. Plant Physiol 155, 2–18.

2 Noctor G, Queval G, Mhamdi A, Chaouch S & Foyer CH (2011) Glutathione. Arabidopsis Book 2011, 1–32.

3 Noctor G, Mhamdi A, Chaouch S, Han Y, Neukermans J, Marquez-Garcia B, Queval G & Foyer CH (2012) Glutathione in plants: an integrated overview. Plant Cell Environ 35, 454–484.

4 Foyer CH & Noctor G (2005) Redox homeostasis and antioxidant signaling: A metabolic interface between stress perception and physiological responses. Plant Cell 17, 1866–1875.

5 Smith IK, Vierheller TL & Thorne CA (1989) Properties and functions of glutathione reductase in plants. Physiol Plant 77, 449–456.

6 Cassier-Chauvat C, Marceau F, Farci S, Ouchane S & Chauvat F (2023) The glutathione system: A journey from cyanobacteria to higher eukaryotes. Antioxidants 2023, *Vol* 12*, Page* 1199 **12**, 1199.

7 Deshpande AA, Bhatia M, Laxman S & Bachhawat AK (2017) Thiol trapping and metabolic redistribution of sulfur metabolites enable cells to overcome cysteine overload. Microbial Cell 4, 112.

8 Ohkama-Ohtsu N & Wasaki J (2010) Recent progress in plant nutrition research: Cross-talk between nutrients, plant physiology and soil microorganisms. Plant Cell Physiol 51, 1255– 1264.

9 Ito T & Ohkama-Ohtsu N (2023) Degradation of glutathione and glutathione conjugates in plants. J Exp Bot 74, 3313–3327.

10 Geu-Flores F, Nielsen MT, Nafisi M, Møldrup ME, Olsen CE, Motawia MS & Halkier BA (2009) Glucosinolate engineering identifies a γ-glutamyl peptidase. Nature Chemical Biology 2009 5:8 5, 575–577.

11 Geu-Flores F, Moldrup ME, Böttcher C, Olsen CE, Scheel D & Halkier BA (2011) Cytosolic γ-glutamyl peptidases process glutathione conjugates in the biosynthesis of glucosinolates and camalexin in *Arabidopsis*. Plant Cell 23, 2456–2469.

12 Ito T, Kitaiwa T, Nishizono K, Umahashi M, Miyaji S, Agake S, Kuwahara K, Yokoyama T, Fushinobu S, Maruyama-Nakashita A, Sugiyama R, Sato M, Inaba J, Hirai MY & Ohkama-Ohtsu N (2022) Glutathione degradation activity of γ-glutamyl peptidase 1 manifests its dual roles in primary and secondary sulfur metabolism in *Arabidopsis*. The Plant Journal 111, 1626–1642.

13 Miyaji S, Ito T, Kitaiwa T, Nishizono K, Agake SI, Harata H, Aoyama H, Umahashi M, Sato M, Inaba J, Fushinobu S, Yokoyama T, Maruyama-Nakashita A, Hirai MY & Ohkama-Ohtsu N (2024) *N*^2^-Acetylornithine deacetylase functions as a Cys-Gly dipeptidase in the cytosolic glutathione degradation pathway in *Arabidopsis thaliana*. The Plant Journal 118, 1603–1618.

14 Rawlings ND, Barrett AJ, Thomas PD, Huang X, Bateman A & Finn RD (2018) The MEROPS database of proteolytic enzymes, their substrates and inhibitors in 2017 and a comparison with peptidases in the PANTHER database. Nucleic Acids Res 46, D624–D632.

15 Krissinel E & Henrick K (2007) Inference of macromolecular assemblies from crystalline state. J Mol Biol 372, 774–797.

16 Chave KJ, Auger IE, Galivan J & Ryan TJ (2000) Molecular modeling and site-directed mutagenesis define the catalytic motif in human γ-glutamyl hydrolase. Journal of Biological Chemistry 275, 40365–40370.

17. Li H, Ryan TJ, Chave KJ & Van Roey P (2002) Three-dimensional Structure of Human γ-Glutamyl Hydrolase: A CLASS I GLUTAMINE AMIDOTRANSFERASE ADAPTED FOR A COMPLEX SUBSTRATE. Journal of Biological Chemistry 277, 24522–24529.

18 Schechter I & Berger A (1967) On the size of the active site in proteases. I. Papain. Biochem Biophys Res Commun 27, 157–162.

19 Berg JM, Gatto GJ, Hines J, Tymoczko JL & Stryer L (2023) 6. Enzyme Catalytic Strategies. In Biochemistry 10th Edition, pp. 179–209. W. H. Freeman, San Francisco.

20 Thoden JB, Miran SG, Phillips JC, Howard AJ, Raushel FM & Holden HM (1998) Carbamoyl phosphate synthetase: Caught in the act of glutamine hydrolysis. Biochemistry 37, 8825–8831.

21 Holm L (2020) DALI and the persistence of protein shape. Protein Sci 29, 128–140.

22 Schwarzenbacher R, Deacon AM, Jaroszewski L, Brinen LS, Canaves JM, Dai X, Elsliger MA, Floyd R, Godzik A, Grittini C, Grzechnik SK, Klock HE, Koesema E, Kovarik JS, Kreusch A, Kuhn P, Lesley SA, McMullan D, McPhillips TM, Miller MD, Morse A, Moy K, Nelson MS, Ouyang J, Page R, Robb A, Quijano K, Spraggon G, Stevens RC, Van Den Bedem H, Velasquez J, Vincent J, Von Delft F, Wang X, West B, Wolf G, Hodgson KO, Wooley J & Wilson IA (2004) Crystal structure of a putative glutamine amido transferase (TM1158) from *Thermotoga maritima* at 1.7 Å resolution. *Proteins: Structure*, Function, and Bioinformatics 54, 801–805.

23 Chen Y, Jia H, Zhang J, Liang Y, Liu R, Zhang Q & Bartlam M (2021) Structure and mechanism of the γ-glutamyl-γ-aminobutyrate hydrolase SpuA from *Pseudomonas aeruginosa*. Acta Crystallogr D Struct Biol 77, 1305–1316.

24 Knöchel T, Ivens A, Hester G, Gonzalez A, Bauerle R, Wilmanns M, Kirschner K & Jansonius JN (1999) The crystal structure of anthranilate synthase from *Sulfolobus solfataricus*: Functional implications. Proc Natl Acad Sci U S A 96, 9479–9484.

25 Spraggon G, Kim C, Nguyen-Huu X, Yee MC, Yanofsky C & Mills SE (2001) The structures of anthranilate synthase of *Serratia marcescens* crystallized in the presence of (i) its substrates, chorismate and glutamine, and a product glutamate, and (ii) its end-product inhibitor, L-tryptophan. Proc Natl Acad Sci U S A 98, 6021–6026.

26 Zubieta C, Krishna SS, McMullan D, Miller MD, Abdubek P, Agarwalla S, Ambing E, Astakhova T, Axelrod HL, Carlton D, Chiu HJ, Clayton T, Deller M, DiDonato M, Duan L, Elsliger MA, Grzechnik SK, Hale J, Hampton E, Gye WH, Haugen J, Jaroszewski L, Jin KK, Klock HE, Knuth MW, Koesema E, Kumar A, Marciano D, Morse AT, Nigoghossian E, Oommachen S, Reyes R, Rife CL, Van Den Bedem H, Weekes D, White A, Xu Q, Hodgson KO, Wooley J, Deacon AM, Godzik A, Lesley SA & Wilson IA (2007) Crystal structure of homoserine *O*-succinyltransferase from *Bacillus cereus* at 2.4 Å resolution. *Proteins: Structure*, Function, and Bioinformatics 68, 999–1005.

27 Sagong HY, Lee D, Kim IK & Kim KJ (2022) Rational engineering of homoserine *O*-succinyltransferase from *Escherichia coli* for reduced feedback inhibition by methionine. J Agric Food Chem 70, 1571–1578.

28 Chandonia JM, Guan L, Lin S, Yu C, Fox NK & Brenner SE (2022) SCOPe: improvements to the structural classification of proteins – extended database to facilitate variant interpretation and machine learning. Nucleic Acids Res 50, D553–D559.

29 Stark AA, Zeiger E & Pagano DA (1993) Glutathione metabolism by γ-glutamyltranspeptidase leads to lipid peroxidation: characterization of the system and relevance to hepatocarcinogenesis. Carcinogenesis 14, 183–189.

30 Levine A, Tenhaken R, Dixon R & Lamb C (1994) H_2_O_2_ from the oxidative burst orchestrates the plant hypersensitive disease resistance response. Cell 79, 583–593.

31 Desikan R, Reynolds A, Hancock JT & Neill SJ (1998) Harpin and hydrogen peroxide both initiate programmed cell death but have differential effects on defence gene expression in *Arabidopsis* suspension cultures. Biochemical Journal 330, 115–120.

32 Desikan R, A.-H.-Mackerness S, Hancock JT & Neill SJ (2001) Regulation of the *Arabidopsis* transcriptome by oxidative stress. Plant Physiol 127, 159–172.

33 Higdon J V., Delage B, Williams DE & Dashwood RH (2007) Cruciferous vegetables and human cancer risk: epidemiologic evidence and mechanistic basis. Pharmacol Res 55, 224–236.

34 Hayes JD, Kelleher MO & Eggleston IM (2008) The cancer chemopreventive actions of phytochemicals derived from glucosinolates. Eur J Nutr 47, 73–88.

35 Yamashita K, Hirata K & Yamamoto M (2018) KAMO: towards automated data processing for microcrystals. Acta Crystallogr D Struct Biol 74, 441–449.

36 Kabsch W (2010) XDS. Acta Crystallogr Sect D-Struct Biol 66, 125–132.

37 Winter G, Waterman DG, Parkhurst JM, Brewster AS, Gildea RJ, Gerstel M, Fuentes-Montero L, Vollmar M, Michels-Clark T, Young ID, Sauter NK & Evans G (2018) DIALS: implementation and evaluation of a new integration package. urn:issn:2059-7983 74, 85– 97.

38 Evans PR & Murshudov GN (2013) How good are my data and what is the resolution? Acta Crystallogr Sect D-Struct Biol 69, 1204–1214.

39 McCoy AJ, Grosse-Kunstleve RW, Adams PD, Winn MD, Storoni LC & Read RJ (2007) Phaser crystallographic software. J Appl Crystallogr 40, 658–674.

40 Mirdita M, Schütze K, Moriwaki Y, Heo L, Ovchinnikov S & Steinegger M (2022) ColabFold: making protein folding accessible to all. Nature Methods 2022 19:6 19, 679– 682.

41 Emsley P, Lohkamp B, Scott WG & Cowtan K (2010) Features and development of Coot. Acta Crystallogr Sect D-Struct Biol 66, 486–501.

42 Murshudov GN, Skubák P, Lebedev AA, Pannu NS, Steiner RA, Nicholls RA, Winn MD, Long F & Vagin AA (2011) REFMAC5 for the refinement of macromolecular crystal structures. Acta Crystallogr D Biol Crystallogr 67, 355–367.

43 Liebschner D, Afonine P V., Moriarty NW, Poon BK, Sobolev O V., Terwilliger TC & Adams PD (2017) Polder maps: Improving OMIT maps by excluding bulk solvent. Acta Crystallogr D Struct Biol 73, 148–157.

44 Pettersen EFF, Goddard TDD, Huang CCC, Couch GSS, Greenblatt DMM, Meng ECC & Ferrin TEE (2004) UCSF Chimera - A visualization system for exploratory research and analysis. J Comput Chem 25, 1605–1612.

45 Pettersen EF, Goddard TD, Huang CC, Meng EC, Couch GS, Croll TI, Morris JH & Ferrin TE (2021) UCSF ChimeraX: Structure visualization for researchers, educators, and developers. Protein Science 30, 70–82.

46 Micsonai A, Moussong É, Wien F, Boros E, Vadászi H, Murvai N, Lee YH, Molnár T, Réfrégiers M, Goto Y, Tantos Á & Kardos J (2022) BeStSel: webserver for secondary structure and fold prediction for protein CD spectroscopy. Nucleic Acids Res 50, W90– W98.

